# Deciphering a hexameric protein complex with Angstrom optical resolution

**DOI:** 10.1101/2021.11.18.468928

**Authors:** Hisham Mazal, Franz-Ferdinand Wieser, Vahid Sandoghdar

**Affiliations:** Max Planck Institute for the Science of Light, 91058 Erlangen, Germany; Max-Planck-Zentrum für Physik und Medizin, 91058 Erlangen, Germany; Friedrich-Alexander University of Erlangen-Nürnberg, 91058 Erlangen, Germany

**Keywords:** Cryogenic super-resolution, correlative imaging, protein structure and assembly, fluorescence

## Abstract

Cryogenic optical localization in three dimensions (COLD) was recently shown to resolve up to four binding sites on a single protein. However, because COLD relies on intensity fluctuations that result from the blinking behavior of fluorophores, it is limited to cases, where individual emitters show different brightness. This significantly lowers the measurement yield. To extend the number of resolved sites as well as the measurement yield, we employ partial labeling and combine it with polarization encoding in order to identify single fluorophores during their stochastic blinking. We then use a particle classification scheme to identify and resolve heterogenous subsets and combine them to reconstruct the three-dimensional arrangement of large molecular complexes. We showcase this method (polarCOLD) by resolving the trimer arrangement of proliferating cell nuclear antigen (PCNA) and the hexamer geometry of Caseinolytic Peptidase B (ClpB) of *Thermus thermophilus* in its quaternary structure, both with Angstrom resolution. The combination of polarCOLD and single-particle cryogenic electron microscopy (cryoEM) promises to provide crucial insight into intrinsic, environmental and dynamic heterogeneities of biomolecular structures. Furthermore, our approach is fully compatible with fluorescent protein labeling and can, thus, be used in a wide range of studies in cell and membrane biology.

**Significance statement:** Fluorescence super-resolution microscopy has witnessed many clever innovations in the last two decades. Here, we advance the frontiers of this field of research by combining partial labeling and 2D image classification schemes with polarization-encoded single-molecule localization at liquid helium temperature to reach Angstrom resolution in three dimensions. We demonstrate the performance of the method by applying it to trimer and hexamer protein complexes. Our approach holds great promise for examining membrane protein structural assemblies and conformations in challenging native environments. The methodology closes the gap between electron and optical microscopy and offers an ideal ground for correlating the two modalities at the single-particle level. Indeed, correlative light and electron microscopy is an emerging technique that will provide new insight into cell biology.

## Introduction

Proteins and their various assemblies are among the main constituents of all living systems and govern every aspect of cellular physiology in both healthy and disease states (1-4). These biomolecular structures adopt sophisticated three-dimensional (3D) configurations during their multifaceted conformational changes. A full understanding of their spatial arrangements and associated heterogeneous configurations is crucial for elucidating their molecular mechanisms, helps guide the engineering of new proteins, and is a great asset for drug discovery (5). Indeed, since the pioneering work of Perutz on protein crystals (6), a variety of techniques such as X-ray crystallography (7) and nuclear magnetic resonance (NMR) spectroscopy (8) have been explored for gaining insight in protein structure and function. Advances in sample preparation, detector technology, and image processing based on single-particle analysis have also ushered in atomic resolution in cryogenic electron microscopy (cryoEM) studies of protein structure (9, 10). The inherent resolution of this method is highly desirable, but lack of specific labeling makes it challenging to identify small variations such as structural inhomogeneities. Fluorescence microscopy, on the other hand, draws its success from an exquisite specificity in labeling, but has traditionally suffered from a limited resolution.

The recent advent of super-resolution (SR) fluorescence microscopy has opened fully new avenues for studying sub-cellular organization and is on the way to become a workhorse for biological studies (11-13). However, conventional SR microscopy performed at room temperature is still not considered as a contestant in the arena of structural biology, where Angstrom-level information about the molecular architecture of proteins and protein complexes is sought after. To push the limit of fluorescence microscopy, one can perform measurements under cryogenic conditions (14-25). In addition to slowing down photochemistry, which allows each fluorophore to emit several orders of magnitude more photons than at room temperature (23-25), a key advantage of cryogenic temperatures is in offering superior sample preservation and high stability for Angstrom-scale structural studies. In one implementation, cryogenic optical localization in 3D (COLD) was introduced, where the stochastic intensity blinking of organic dyes gave access to the positions of up to four labeling sites on a single protein (21). Recently, we employed a more robust protocol to identify individual fluorophores by exploiting the polarization of the fluorescence light dictated by the fluorophore orientation (14). This latter method was validated by measuring single distances on one-dimensional DNA nanorulers. In our current study, we introduce polarCOLD, which exploits polarization encoding for resolving several fluorophores in 3D. Importantly, we show that the distances and arrangements of protein complexes can be determined by combining images recorded from under-sampled structures. To demonstrate this, we first resolve three fluorophores on a trimer protein complex with Angstrom resolution. Next, we use partial labeling, a supervised particle classification procedure, and the particle symmetry to solve a complete hexameric protein arrangement. We discuss the promise of our methodology for combination with cryoEM.

## Results

### polarCOLD on a protein trimer

Proliferating cell nuclear antigen (PCNA) is a central functional unit in genome repair and replication (26). The structure of this complex protein was solved by X-ray crystallography (27) and more recently by cryoEM (28). These studies have shown that PCNA forms a stable homo-trimer with a pseudo-hexameric shape. The stability and simple configuration of its structure makes PCNA a good model system for benchmarking our imaging methodology (14). To study Human PCNA with polarCOLD, we first fully labeled it via a His-tag linker on the N-terminal side of each subunit of the protein, forming an equilateral triangle (see method section and Fig. S1). PCNA complexes were then embedded in a hydrophilic poly-vinyl alcohol (PVA) matrix at sub-nanomolar concentration and spin-coated on a mirror-enhanced substrate (14). The resulting density corresponds to fewer than one protein per μm^2^ on average so that individual proteins can be easily resolved in diffraction-limited imaging. The samples were immediately imaged in our custom-built microscope (see method section).

Figure 1a shows a schematic of the imaging setup. A liquid helium cryostat houses the sample stage as well as a scannable microscope objective (14, 21). A polarizing beam splitter in the detection path allows us to determine the orientation of dipole-like emitters projected onto the angular interval θ ϵ [0°, 90°] in the imaging plane. The high photostability of the fluorophores at T=4K allows us to collect on average 260 photons per frame from a single fluorophore with a total number of registered photons exceeding 10^6^ after 50,000 frames (see Fig. S2). Figure 1b-c (blue trace) displays two examples in which three different polarization states recur at various times. The long photo-blinking off-times (see also Fig. S2e) allow one to identify each of the three fluorophores on a given protein complex separately (21). More insight into polarization trace processing can be found in Fig. S3 of the SI. To resolve the polarization histograms in cases where they partially overlap (see, e.g., Fig. 1c), we fit the data using an algorithm that combines unsupervised statistical learning tools with change-point detection in a model-independent manner (29). As illustrated by the red traces in Fig. 1b,c, we can robustly identify the polarization states over time and hence assign the signal in each frame to a single fluorophore. The maximum number of resolvable polarization states per protein is currently limited to fewer than 5 fluorophores, dictated by the width of the polarization histogram (Fig. S2a). We remark that the reliability of this procedure can be arbitrarily high if one sacrifices measurement yield because one can simply exclude the cases for which the expected number of polarization states is not well resolved (see Table S1 for the statistics of this analysis).

**Figure 1.**
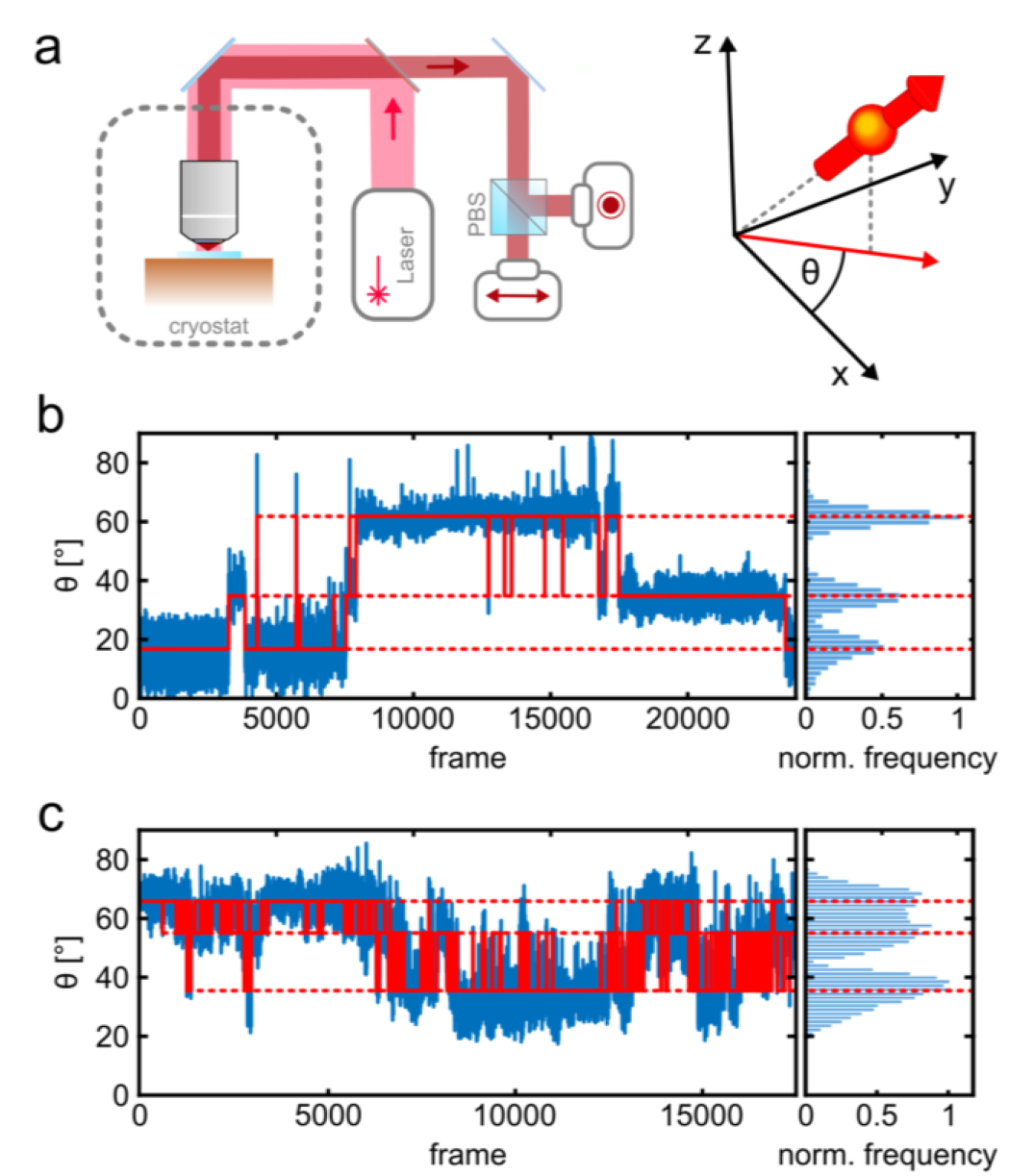
Photo-physics and co-localization of fluorophores at cryogenic temperatures. (a) Schematics of the cryogenic optical microscope. Polarization-resolved detection allows for direct measurement of the in-plane dipole moment of fluorophores. Here we use a circularly polarized light from a laser at λ = 635 nm. A polarizing beam splitter in the detection path allows one to resolve the polarization state of each individual molecule. (b, c) Exemplary polarization time traces of two single proteins. (b) demonstrates a case of a well-separated polarization states, whereas (c) displays a case with smaller separations between polarization states. Blue traces present the experimental polarization values for each frame, and the red lines show the polarization determined by the algorithm (29). The blinking kinetics are exceptionally slow with on/off times in the range of seconds to minutes.

Having identified the individual fluorophores, we generate super-resolved images by clustering the respective coordinates and taking their averages. The top and bottom rows in Fig. 2a display a selection of the measured and simulated 2D projection maps. To quantify their similarity, we computed a correlation score ranging from 0 to 1. We obtained 2D correlation scores of 0.92 or higher, representing nearly perfect agreement. To obtain a 3D model from our 2D localization maps, we use a single-particle reconstruction algorithm (21, 30) (see method section and Fig. S4 for complete data set). Figure 2b shows that the reconstructed fluorophore volumes (red spheroids) agree well with the crystal structure of the PCNA protein (PDB: 1AXC) containing 3 identical subunits in an equilateral triangle. The slight asymmetry and deviation from the actual crystal structure can in part be attributed to the uncertainty introduced by the dye linker, 6-histidine linker and possibly the restricted rotational mobility of the dye itself, resulting in a minor localization bias. Indeed, by taking the dye linker into account and calculating the accessible volume (31), we found that our 3D reconstructed volumes correlate very well (0.96 correlation score) with the simulated accessible volume (see Fig. 2b).

**Figure 2.**
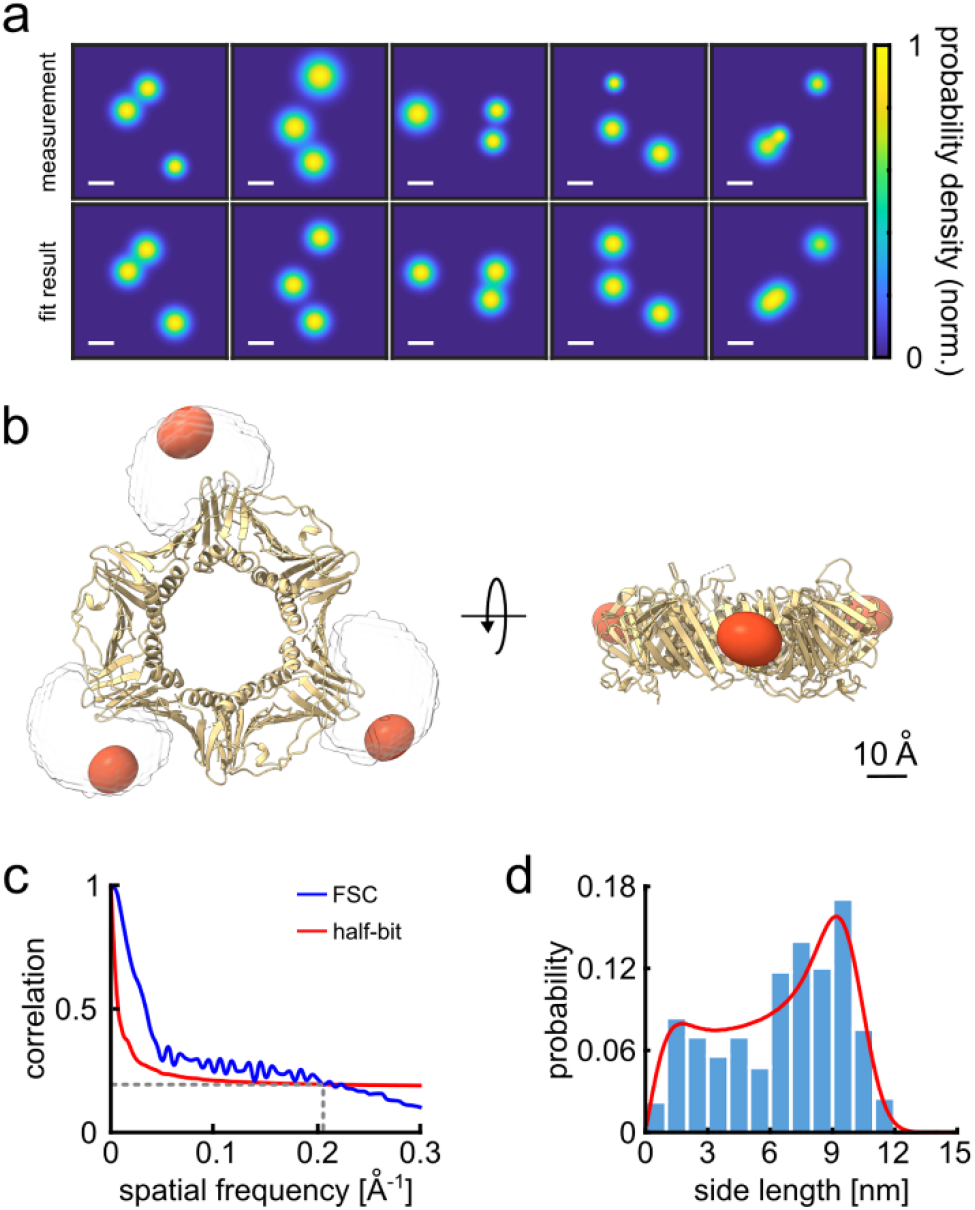
3D reconstruction of the PCNA protein trimer. (a) Experimentally-obtained super-resolved 2D images (top row) of single proteins and simulated images based on the crystal structure (bottom row). The color code represents the occupation probability determined by the localization precision for each fluorophore. The localization precision in the simulated data was normalized. Scale bar is 3 nm. (b) Overlay of the crystal structure of human PCNA with the reconstructed fluorophore volumes shown as red spheroids (see online Animation 1). The transparent white clouds represent the accessible volume of the dye linker attached to the N-terminal side of the protein, calculated using the parameter of ATTO647N as provided in Ref. (31). By fitting the reconstructed 3D volumes obtained from polarCOLD into the theoretical accessible volumes of the dyes, we find a correlation score of 0.96, indicating a correct 3D reconstruction. (c) The Fourier shell correlation (FSC, blue curve) of the two half data sets gives a resolution of 4.9 Å based on the half-bit criterion (red curve). (d) Distribution of the projected side lengths (blue) obtained from the localized positions shown in (b). The model fit (red) takes the finite localization uncertainty and the random particle orientation into account, resulting in 9.9 ± 0.6 nm.

We also verified the robustness of our assignment procedure by performing random assignment of frames, which resulted in single, unresolved spots (see method section and Fig. S3). To evaluate the resolution of our 3D reconstructed volumes, we used the well-established method of Fourier shell correlation (FSC) (32). Here, we divide the 2D image data set into two randomly chosen groups and then determine their 3D reconstruction separately. Then we assess the cross-correlation (similarity) between the two 3D volumes in Fourier space as a function of spatial frequency. The overall resolution of a 3D reconstructed volume is thus obtained by finding the maximum spatial frequency corresponding to a correlation above a specific threshold value. Here, we used the half-bit criteria, which is a standard threshold curve used in single particle cryoEM. As shown in Fig. 2c, the intersection of this curve (red) with the FSC curve (blue) indicates at which spatial frequency we have collected a sufficient amount of information in order to interpret the 3D reconstructed volumes accurately (32). We find a remarkable resolution of 4.9 Å. We further quantified the size of the protein-dye conjugate via the pair-wise distances between the localized sites on each particle. The histogram in Fig. 2d plots the distribution of the side lengths of the projected triangles and is well-described by a fit that considers the localization uncertainty as well as the random particle orientation (14). We determine a side length of 9.9 ± 0.6 nm in excellent agreement with the expected value. The uncertainty was determined via bootstrapping and is consistent with the resolution obtained from the FSC curve. We note that the high signal-to-noise ratio (SNR) of the method (see, e.g, Fig. 2a) produces good results from a total of only 119 particles (see Table S1 for overall statistics), which is two to three orders of magnitude lower than the number required for typical cryoEM measurements (33).

### Resolving a hexameric protein complex using partial labeling

The trimer structure discussed above involves only a single dye-dye distance. We now turn to resolving an example of more complex higher-order protein structures. Considering that a limited number of fluorophores can be resolved via stochastic blinking (Fig. S2a), we pursued a strategy of partial labeling of the sites of interest on a given individual protein complex. The piece-wise information, which involves various fluorophore arrangements and distances is then assembled to solve for the full architecture using prior knowledge of the symmetry. This concept has been successfully used to build structural models in NMR (34) and more recently in SR microscopy (35, 36). To demonstrate this technique, we examine the homo-hexamer Caseinolytic Peptidase B (ClpB) of *Thermus thermophilus* in its quaternary structure (37, 38) (Fig. S5a). ClpB is a molecular machine that rescues proteins from aggregation within cells (39), and its structure was shown to be very stable in the presence of ATP at low concentrations (40). Here, we labeled the M-domain of the protein at residue S428C such that the distance between two adjacent labeling sites is expected to be 9 nm after accounting for the dye linker (Fig. S5a). We deliberately reduced the labeling efficiency to 50% in order to allow for 1/3 of the particles to carry three fluorophores as estimated by the binomial distribution (Fig. S5b).

Labeled ClpB complexes were imaged in the same fashion as before in the presence of 2 mM ATP to stabilize the protein assembly. We obtained about 510 photons per frame on average and a significantly higher total number of photons obtained after 100,000 frames since 95% of the molecules survived until the end of the acquisition interval (Fig. S2f-i). The photoblinking behavior was similar to that observed in PCNA with a slight increase in the average off-on ratio, allowing robust fluorophore assignment. Hence, we fitted our polarization traces as described previously and selected only those cases that contained three fluorophores. Particles carrying three labels inherently fall into three distinct classes (I, II and III) with multiple pair-wise distances of approximately 9, 15 and 18 nm (see Fig. S5a).

To analyze the resulting triangle images, we used a supervised classification scheme (41) to assign each 2D image to one of the three classes, taking the angle of its plane into account. Here, we simulated large data sets of 2D projections for each class and applied a template matching procedure based on the 2D cross-correlation between an experimental image and the simulated images. A control based on simulated ground truth images obtained from the crystal structure of ClpB showed an accuracy of 97% in template matching. Figure 3a displays examples of the 2D super-resolved images of each class (see Fig. S6 for more examples of different projections). We, thus, assigned each image to the class that yielded the highest correlation score. Images that were fitted to all classes with a difference below 10% (estimated from a simulation analysis) in the score were excluded from further analysis in order to avoid smearing (see Table S1 for all statistics).

**Figure 3.**
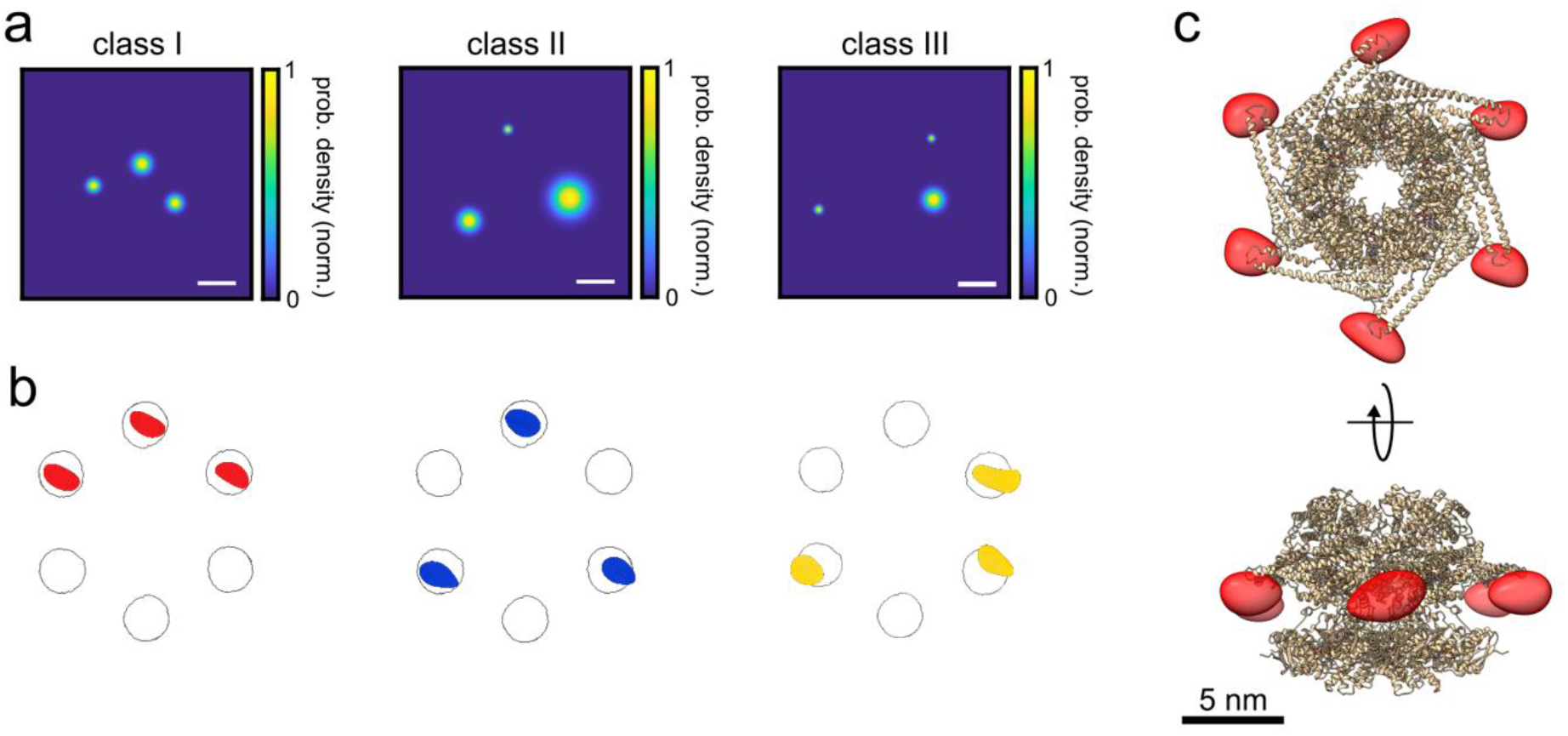
3D reconstruction of the ClpB hexamer. (a) Top view of super-resolved 2D images for class I, II and III, as obtained from single-particle classification procedure. Scale bar is 3 nm. (b) Result of single-particle classification and averaging. The reconstructed 3D volume of each class nicely sits in the simulated accessible volume of the fluorophores (grey spheres). Red, blue and yellow spheres represent class I, II and III with correlation values of 0.98, 0.96 and 0.86, respectively. (c) 3D reconstruction of the complete hexamer obtained from merging the 3D volumes (red spheroid) of the three classes. Top figure shows the top view of the reconstructed 3D volume of the hexamer shape, and the bottom figure shows its 90° rotation (see Animation 2 and 3). Crystal structure of ClpB is shown as a cartoon in gold (PDB: 1QVR) (37, 38).

Next, we picked 2D images of each class with a localization precision better 3 nm (Fig. S5c) and correlation score better than 0.9 and calculated their respective 3D structures as described previously for PCNA. Overall, we obtained 232, 100, and 135 particles for class 1, 2 and 3, respectively, reaching a yield of 7,46% for all the particles detected with three polarization states (see Table S1). Remarkably, as illustrated in Fig. 3b, the reconstructed 3D volumes fit very well to the probable locations of the dye on the protein structure given by the accessible volumes (circles) on each protomer after taking the dye linker into account. The correlation scores between the reconstructed 3D volumes and the accessible volumes of the dyes were 0.98, 0.96 and 0.86 for class I, II and III, respectively. As a control, we also performed a reconstruction of unclassified 2D images, and confirmed that no structure was identified (Fig. S7). Again, the FSC curves of the volumes (see Fig. S5d-f) suggest an exquisite resolution of 4.0, 7.9 and 6.4 Å, for class I, II and III, respectively.

Following the successful assignment of the 3 classes, we merged them as shown in Fig. 3c to obtain the complete 3D shape of the hexamer structure. We note that classification approaches reported recently in SR fluorescent microscopy (36, 42) have focused on samples where the particle orientation is either naturally or artificially fixed in a given direction, e.g. a top view. Our results present the first study on classifying heterogenous configurations on the single-protein level including all possible orientations.

## Discussion

polarCOLD can be further improved and extended to other modalities, e.g., via combination with genetic labeling (15, 43-45) or by exploiting the narrow absorption and emission spectra at low temperatures for co-localization of larger numbers of fluorescent markers. However, the currently achieved SNR and Angstrom resolution of cryogenic light microscopy is already capable of providing pivotal solutions for quantitative structural analysis of small proteins as well as large protein complexes and aggregates. A specially promising line of study concerns the conformation of membrane proteins in their native environment. Indeed, an estimated 20% of the human genome encodes membrane proteins and many of them are potential drug targets (46). The structure and active configuration of many proteins and protein complexes, however, remain out of reach because it is very challenging to crystallize or resolve them in their crowded surrounding (47). In particular, heterogeneities caused by mobile domains and intrinsic distributions in assemblies cannot be preserved and classified in a robust fashion so that they lead to a smearing effect in the final reconstruction in EM studies (48-53). A future direction is, thus, to prepare and vitrify biological samples via rapid freezing (54) or high-pressure freezing (15, 55) in order to exploit the high spatial resolution, sensitivity and specificity of polarCOLD for identifying and determining the positions and orientations of individual particles of interest in low-contrast EM micrographs.

The combination of cryogenic light microscopy with cryoEM has attracted attention for cellular and tissue imaging although the existing reports do not surpass a resolution of about 10 nm (15, 16, 56-69). A very exciting prospect is now to combine polarCOLD and cryoEM on the single-particle level. As illustrated in Fig. S8, by labeling a functional domain within a single molecule with two or three fluorophores, it should be possible to classify the particles based on conformational changes as indicated by their intermolecular distances. Similarly, by labeling two different protomers or functional domains within a protein complex, it will be possible to group the particles based on intramolecular distances that result from conformational changes between subunits. In addition, while we currently exploit the prior symmetry knowledge of the structures under study, we plan to explore unsupervised classification schemes for identifying and classifying particles in an unknown sample. Altogether, these approaches will help to overcome various challenges connected to low purification yields and large backgrounds from cell membranes, paving the way to studying samples containing a heterogeneous distribution of similar structures and mapping their energy landscapes.

## Materials and Methods

### Protein labeling

His-tagged human PCNA protein was purchased from Sigma Aldrich (catalogue number SRP5117), at a concentration of 6.6 μM. The protein was specifically labeled via the histidine linker on the N-terminal side of the protein with the dye Atto647N containing a Ni-NTA functional group which was also purchased from Sigma Aldrich (catalogue number 02175-250UG-F). First, the protein was desalted to a labeling buffer (25 mM HEPES, 25 mM KCl, pH 7.8) using a 7K MWCO Zeba desalting column (ThermoFischer, cat. 89882) and then reacted with dyes at a ratio of 1:4 for 2 hr at RT. The protein was then desalted from the excess of dyes using the same desalting column. The labeling efficiency was estimated using an absorption spectrometer (Nanodrop 2000, ThermoFischer) confirming ~100% labeling efficiency. SDS-page and native gel indicated that the protein is indeed assembled and labeled. N-terminal His-tagged ClpB protein, mutated at residue 428 from alanine to cysteine, was purified as described in a previous publication (40). The labeling of the single cysteine was done following the same procedure as describe above with Atto647 maleimide as a specific labeling agent of the cysteine. Absorption spectroscopy, SDS-page and native gel indicate complete assembly of the protein, see previous publication (40).

### Sample preparation

Both protein samples were prepared in a similar way, and all the samples were prepared freshly on the same day. Proteins were diluted to a stock solution of 50 nM, in 25 mM HEPES, 25 mM KCl, 10 mM MgCl_2_, 0.5 mM TCEP at pH 7.8 (working buffer). In the case of ClpB 2mM ATP was added to stabilize protein assembly. The stock solution of poly-vinyl alcohol (PVA) was prepared as follows: 15 μl of 8% PVA (0.3% final concentration) was diluted into 345 μl working buffer containing 1 mM Trolox and 2 mM ATP in the case of ClpB. Then 2 μl of the protein was mixed with the 360 μl PVA solution and filtered with a 100 nm spin filter (Whatman Anotop-10). 5 μl of this mixed solution was then spin coated (30 s at 1000 rpm followed by 3000 rpm for 60 s) onto a plasma-cleaned mirror-enhanced substrate, prepared in house, (14), and immediately loaded into our custom-built cryogenic microscope.

### Experimental setup

All experiments were performed in a cryogenic microscope that is built around a Janis ST-500 flow cryostat and operates at liquid helium temperature. Samples are loaded onto a cold finger and imaged by a 0.95 NA objective (Olympus MPLAPO 100x), which is mounted in vacuum, onto two separate EMCCD cameras (Andor iXon) in a polarization-resolved configuration. A more detailed description of the optical setup can be found in Ref. (14). The laser intensity used in all experiments was set to ~1 kW/cm^2^ and images were recorded at 70 Hz. For each field of view, we collected a total of 50,000 or 100,000 frames for PCNA and ClpB, respectively. Our field of view was 211×313 pixels with a pixel size of 190 nm.

### Image analysis

We analyzed raw image stacks from two polarization channels with custom-written MATLAB software. Briefly, we perform dual-channel localization with a Poisson-weighted Gaussian maximum-likelihood estimator. Localized coordinates are drift corrected in a two-step process and registered to nanometer accuracy via a nonlinear alignment. Polarization time traces are then analyzed (see Fig. S3) by a routine that is based on the DISC algorithm (29) to find the polarization states. Once all polarization states are identified, the associated coordinates are averaged to generate a 2D super-resolved image. Here in our case, we took the particles which were fitted best to a three polarization states, and used them for further analysis (see Table S1 for statistics).

### Single-molecule 2D image fitting

To fit the 2D maps of the human PCNA, we used the theoretical distances obtained from the structural model of human PCNA (PDB: 1AXC) and then we performed a least-squares fit over rotation angles and translation using a simulated annealing algorithm. Two-dimensional cross-correlation of the experimental and simulated maps yield a score better than 0.92. In the case of the hexamer protein, ClpB, we generated a large space of multiple 2D projection, ~60,000 images for each class, using a rotation matrix over x, y, and z axis with a resolution angle of 10°. The exact distances where evaluated from the real crystal structural model which include the dye linker into account (ClpB model, PDB: 1QVR, and the full assembly model taken from Ref (38)). Then we performed 2D template matching between each experimental data and all the images from the simulated data. Here, we used a 2D cross-correlation algorithm to calculate the similarity index between two images. The correlation score ranges from 0 to 1, where 1 indicates a 100% match. For each experimental image we select the fit with the highest score as the best fit to our data, and to assign it to a class. Images that were fitted to all classes with a difference below 10% in the score were excluded from further analysis to avoid smearing.

### Tomography and 3D reconstruction

Here we selected the particles which were fitted properly to the simulated 2D projections. Namely we selected particles with a cross-correlation score better than 0.9 and with a localization precision better than 3 nm. The 2D maps of the particles were then normalized (maximum and width) to perform the 3D reconstruction. For a full 3D reconstruction, we used the subspaceEM algorithm (30) with default settings and 100 runs. The algorithm was initialized with an elliptical Gaussian placed in the center of the volume.

The 2D maps of the trimer protein, PCNA, were constructed using a grid size of 120×120 with a pixel size of 1.5 Å. In the case of the hexamer protein, the grid size was 200×200 with a pixel size of 1.5 Å. The 3D volumes were then processed, fitted to the crystal structures/maps of the dye location and edited using ChimeraX (70). The Fourier shell correlation curves (FSC), were computed using the freely available FSC server, provided by Protein Data Bank in Europe website (https://www.ebi.ac.uk/pdbe/emdb/validation/fsc/). Here we divide the 2D images data sets into two equally data sets, and calculate their 3D reconstruction similar to as described above. Then we align the two reconstituted volumes and calculate the FSC and detriment the resolution based on half-bit criteria (32).

### Distance fitting

For more quantitative analysis of the pair-wise distances we used a model described in Ref. (14, 21). In short, we convolve the Rician distribution of the distance between two spots with finite localization uncertainty with the projection onto the image plane given by a cosine function.

## Acknowledgments

We thank Tobias Utikal for assistance with cryogenic experiments, reading and commenting on the manuscript and Michelle Küppers for reading and commenting on the manuscript. We thank Simone Ihloff for support with materials, Alexander Gumann for preparing the mirror-enhanced substrates and Jan Renger for help with measuring the PVA film thickness. We are further grateful to Prof. Gilad Haran for the generous gift of ClpB proteins and plasmids. H.M. was supported by the Planning & Budgeting Committee of the Council of Higher Education of Israel. We are grateful to the Max Planck Society for financial support.

## Data and materials availability

Data and codes used in this research are available upon request.

## References

1. D. L. Nelson, M. M. Cox, A. L. Lehninger, Lehninger principles of biochemistry (2017).

2. C. Mavroidis, A. Dubey, M. L. Yarmush, Molecular machines. Annu Rev Biomed Eng 6, 363–395 (2004).

3. M. Schliwa, G. Woehlke, Molecular motors. Nature 422, 759 (2003).

4. B. Alberts, The Cell as a Collection of Protein Machines: Preparing the Next Generation of Molecular Biologists. Cell 92, 291–294 (1998).

5. J.-P. Renaud et al., Cryo-EM in drug discovery: achievements, limitations and prospects. Nature Reviews Drug Discovery 17, 471–492 (2018).

6. A. R. Fersht, From the first protein structures to our current knowledge of protein folding: delights and scepticisms. Nat Rev Mol Cell Biol 9, 650–654 (2008).

7. Y. Shi, A glimpse of structural biology through X-ray crystallography. Cell 159, 995–1014 (2014).

8. V. Kanelis, J. D. Forman-Kay, L. E. Kay, Multidimensional NMR methods for protein structure determination. IUBMB Life 52, 291–302 (2001).

9. T. Nakane et al., Single-particle cryo-EM at atomic resolution. Nature 587, 152–156 (2020).

10. W. Kuhlbrandt, The Resolution Revolution. Science 343, 1443–1444 (2014).

11. M. Lelek et al., Single-molecule localization microscopy. Nature Reviews Methods Primers 1 (2021).

12. S. J. Sahl, S. W. Hell, S. Jakobs, Fluorescence nanoscopy in cell biology. Nature Reviews Molecular Cell Biology 18, 685–701 (2017).

13. S. Weisenburger, V. Sandoghdar, Light microscopy: an ongoing contemporary revolution. Contemporary Physics 56, 123–143 (2015).

14. D. Böning, F.-F. Wieser, V. Sandoghdar, Polarization-Encoded Colocalization Microscopy at Cryogenic Temperatures. ACS Photonics 8, 194–201 (2021).

15. D. P. Hoffman et al., Correlative three-dimensional super-resolution and block-face electron microscopy of whole vitreously frozen cells. Science 367 (2020).

16. P. D. Dahlberg et al., Cryogenic single-molecule fluorescence annotations for electron tomography reveal in situ organization of key proteins in Caulobacter. Proc Natl Acad Sci U S A 117, 13937–13944 (2020).

17. F. Moser et al., Cryo-SOFI enabling low-dose super-resolution correlative light and electron cryo-microscopy. Proc Natl Acad Sci U S A 116, 4804–4809 (2019).

18. L. Wang et al., Solid immersion microscopy images cells under cryogenic conditions with 12 nm resolution. Communications Biology 2(2019).

19. C. N. Hulleman, W. Li, I. Gregor, B. Rieger, J. Enderlein, Photon Yield Enhancement of Red Fluorophores at Cryogenic Temperatures. ChemPhysChem 19, 1774–1780 (2018).

20. X. Xu et al., Ultra-stable super-resolution fluorescence cryo-microscopy for correlative light and electron cryo-microscopy. Science China Life Sciences 61, 1312–1319 (2018).

21. S. Weisenburger et al., Cryogenic optical localization provides 3D protein structure data with Angstrom resolution. Nat Methods 14, 141–144 (2017).

22. T. Furubayashi et al., Three-Dimensional Localization of an Individual Fluorescent Molecule with Angstrom Precision. Journal of the American Chemical Society 139, 8990–8994 (2017).

23. W. X. Li, S. C. Stein, I. Gregor, J. Enderlein, Ultra-stable and versatile widefield cryo-fluorescence microscope for single-molecule localization with sub-nanometer accuracy. Optics Express 23, 3770–3783 (2015).

24. S. Weisenburger et al., Cryogenic Colocalization Microscopy for Nanometer-Distance Measurements. ChemPhysChem 15, 763–770 (2014).

25. S. Weisenburger, B. Jing, A. Renn, V. Sandoghdar, Proc. SPIE 8815(2013).

26. I. Bruck, M. O’Donnell, Genome Biology 2, reviews3001.3001 (2001).

27. R. E. Georgescu et al., Structure of a Sliding Clamp on DNA. Cell 132, 43–54 (2008).

28. C. Madru et al., Structural basis for the increased processivity of D-family DNA polymerases in complex with PCNA. Nature Communications 11(2020).

29. D. S. White, M. P. Goldschen-Ohm, R. H. Goldsmith, B. Chanda, Top-down machine learning approach for high-throughput single-molecule analysis. eLife 9(2020).

30. N. C. Dvornek, F. J. Sigworth, H. D. Tagare, SubspaceEM: A fast maximum-a-posteriori algorithm for cryo-EM single particle reconstruction. Journal of Structural Biology 190, 200–214 (2015).

31. S. Kalinin et al., A toolkit and benchmark study for FRET-restrained high-precision structural modeling. Nat Methods 9, 1218–1225 (2012).

32. M. Van Heel, M. Schatz, Fourier shell correlation threshold criteria. Journal of Structural Biology 151, 250–262 (2005).

33. Y. Cheng, N. Grigorieff, Pawel, T. Walz, A Primer to Single-Particle Cryo-Electron Microscopy. Cell 161, 438–449 (2015).

34. J. Fiaux, E. B. Bertelsen, A. L. Horwich, K. Wüthrich, NMR analysis of a 900K GroEL–GroES complex. Nature 418, 207–211 (2002).

35. H. Heydarian et al., Template-free 2D particle fusion in localization microscopy. Nature Methods 15, 781–784 (2018).

36. J. Molle et al., Towards structural biology with super-resolution microscopy. Nanoscale 10, 16416–16424 (2018).

37. S. Lee et al., The Structure of ClpB. Cell 115, 229–240 (2003).

38. A. V. Diemand, A. N. Lupas, Modeling AAA+ ring complexes from monomeric structures. J Struct Biol 156, 230–243 (2006).

39. S. M. Doyle, O. Genest, S. Wickner, Protein rescue from aggregates by powerful molecular chaperone machines. Nat Rev Mol Cell Biol 14, 617–629 (2013).

40. H. Mazal et al., Tunable microsecond dynamics of an allosteric switch regulate the activity of a AAA+ disaggregation machine. Nat Commun 10, 1438 (2019).

41. H. Gao, M. Valle, M. Ehrenberg, J. Frank, Dynamics of EF-G interaction with the ribosome explored by classification of a heterogeneous cryo-EM dataset. Journal of Structural Biology 147, 283–290 (2004).

42. T. A. P. M. Huijben et al., Detecting structural heterogeneity in single-molecule localization microscopy data. Nature Communications 12(2021).

43. P. D. Dahlberg et al., Identification of PAmKate as a Red Photoactivatable Fluorescent Protein for Cryogenic Super-Resolution Imaging. J Am Chem Soc 140, 12310–12313 (2018).

44. B. Liu et al., Three-dimensional super-resolution protein localization correlated with vitrified cellular context. Scientific Reports 5, 13017 (2015).

45. T. M. H. Creemers, A. J. Lock, V. Subramaniam, T. M. Jovin, S. Volker, Photophysics and optical switching in green fluorescent protein mutants. Proceedings of the National Academy of Sciences 97, 2974–2978 (2000).

46. S. Piccoli, E. Suku, M. Garonzi, A. Giorgetti, Genome-wide Membrane Protein Structure Prediction. Current Genomics 14, 324–329 (2013).

47. Y. Cheng, Membrane protein structural biology in the era of single particle cryo-EM. Curr Opin Struct Biol 52, 58–63 (2018).

48. E. Nwanochie, V. N. Uversky, Structure Determination by Single-Particle Cryo-Electron Microscopy: Only the Sky (and Intrinsic Disorder) is the Limit. Int J Mol Sci 20(2019).

49. M. Serna, Hands on Methods for High Resolution Cryo-Electron Microscopy Structures of Heterogeneous Macromolecular Complexes. Frontiers in Molecular Biosciences 6(2019).

50. S. H. W. Scheres, “Chapter Six - Processing of Structurally Heterogeneous Cryo-EM Data in RELION” in Methods in Enzymology, R. A. Crowther, Ed. (Academic Press, 2016), vol. 579, pp. 125–157.

51. S. H. W. Scheres et al., Disentangling conformational states of macromolecules in 3D-EM through likelihood optimization. Nature Methods 4, 27–29 (2007).

52. E. V. Orlova, H. R. Saibil, Structure determination of macromolecular assemblies by single-particle analysis of cryo-electron micrographs. Current Opinion in Structural Biology 14, 584–590 (2004).

53. G. Papai et al., Structure of SAGA and mechanism of TBP deposition on gene promoters. Nature 577, 711–716 (2020).

54. J. Dubochet, A. W. McDowall, VITRIFICATION OF PURE WATER FOR ELECTRON MICROSCOPY. Journal of Microscopy 124, 3–4 (1981).

55. D. Studer, B. M. Humbel, M. Chiquet, Electron microscopy of high pressure frozen samples: bridging the gap between cellular ultrastructure and atomic resolution. Histochemistry and Cell Biology 130, 877–889 (2008).

56. P. D. Dahlberg, W. E. Moerner, Cryogenic Super-Resolution Fluorescence and Electron Microscopy Correlated at the Nanoscale. Annual Review of Physical Chemistry 72(2021).

57. G.-H. Wu et al., Multi-scale 3D Cryo-Correlative Microscopy for Vitrified Cells. Structure 28, 1231–1237.e1233 (2020).

58. M. W. Tuijtel, A. J. Koster, S. Jakobs, F. G. A. Faas, T. H. Sharp, Correlative cryo super-resolution light and electron microscopy on mammalian cells using fluorescent proteins. Scientific Reports 9(2019).

59. G. Wolff, C. Hagen, K. Grünewald, R. Kaufmann, Towards correlative super-resolution fluorescence and electron cryo-microscopy. Biology of the Cell 108, 245–258 (2016).

60. P. De Boer, J. P. Hoogenboom, B. N. G. Giepmans, Correlated light and electron microscopy: ultrastructure lights up! Nature Methods 12, 503–513 (2015).

61. C. Loussert Fonta, B. M. Humbel, Correlative microscopy. Archives of Biochemistry and Biophysics 581, 98–110 (2015).

62. R. Kaufmann, C. Hagen, K. Grünewald, Fluorescence cryo-microscopy: current challenges and prospects. Current Opinion in Chemical Biology 20, 86–91 (2014).

63. Y.-W. Chang et al., Correlated cryogenic photoactivated localization microscopy and cryo-electron tomography. Nature Methods 11, 737–739 (2014).

64. R. Kaufmann et al., Super-Resolution Microscopy Using Standard Fluorescent Proteins in Intact Cells under Cryo-Conditions. Nano Letters 14, 4171–4175 (2014).

65. F. G. A. Faas et al., Localization of fluorescently labeled structures in frozen-hydrated samples using integrated light electron microscopy. Journal of Structural Biology 181, 283–290 (2013).

66. S. Watanabe et al., Protein localization in electron micrographs using fluorescence nanoscopy. Nature Methods 8, 80–84 (2011).

67. A. A. Mironov, G. V. Beznoussenko, Correlative microscopy: a potent tool for the study of rare or unique cellular and tissue events. Journal of Microscopy 235, 308–321 (2009).

68. A. V. Agronskaia et al., Integrated fluorescence and transmission electron microscopy. Journal of Structural Biology 164, 183–189 (2008).

69. A. Sartori et al., Correlative microscopy: Bridging the gap between fluorescence light microscopy and cryo-electron tomography. Journal of Structural Biology 160, 135–145 (2007).

70. T. D. Goddard et al., UCSF ChimeraX: Meeting modern challenges in visualization and analysis. Protein Science 27, 14–25 (2018).

