## Supplementary Information for "Deciphering a hexameric protein complex with Angstrom optical resolution"

### This PDF file includes:

**Figures: S1 – S8**
**Table S1**

### S1 - Characterization and labeling of human PCNA protein

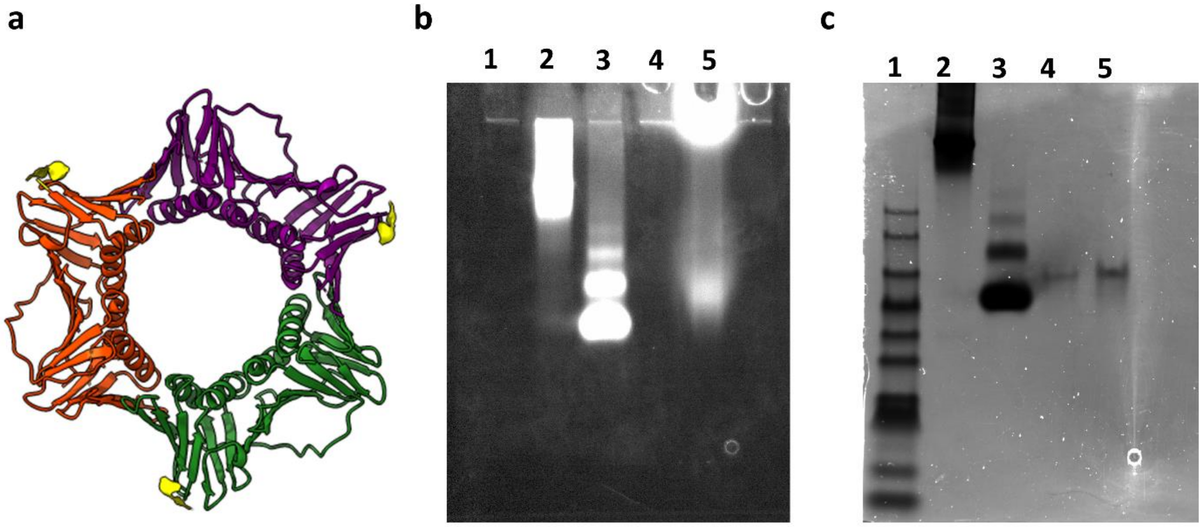


**Figure S1.** (a) Schematics of the human PCNA crystal structure (PDB: 1AXC). Each monomer is presented in a different color (green, red and purple). The location of the N-terminal His-tag is shown in yellow.

Native gel electrophoresis (4–15% Mini-PROTEAN TGX Precast Protein Gels) ran in 25 mM Tris and 192 mM glycine, pH 8, at 60 V for 3 hrs. The gel was imaged with a ChemiDoc XRS system at 647 nm (b) and was then stained with Coomassie Blue (c). Lane 1 is the protein ladder PM2500 (SMOBIO), lane 2 is thyroglobulin from bovine serum, labeled with ATTO647-NHS, lane 3 is Bovine serum albumin protein (BSA) labeled with ATTO647N-NHS, lane 4 is the non-labeled human PCNA, lane 5 is human PCNA labeled with ATTO647N-NTA. Lanes 4 and 5 of human PCNA in the gel show only one band indicating a homogenous and pure solution of the assembled proteins. Also, this result shows that the labeled human PCNA ran similar to the non-labeled one, indicating that the dye did not affect the protein.

### S2 - Fluorophore identification using polarization-resolved imaging and photophysics characterization

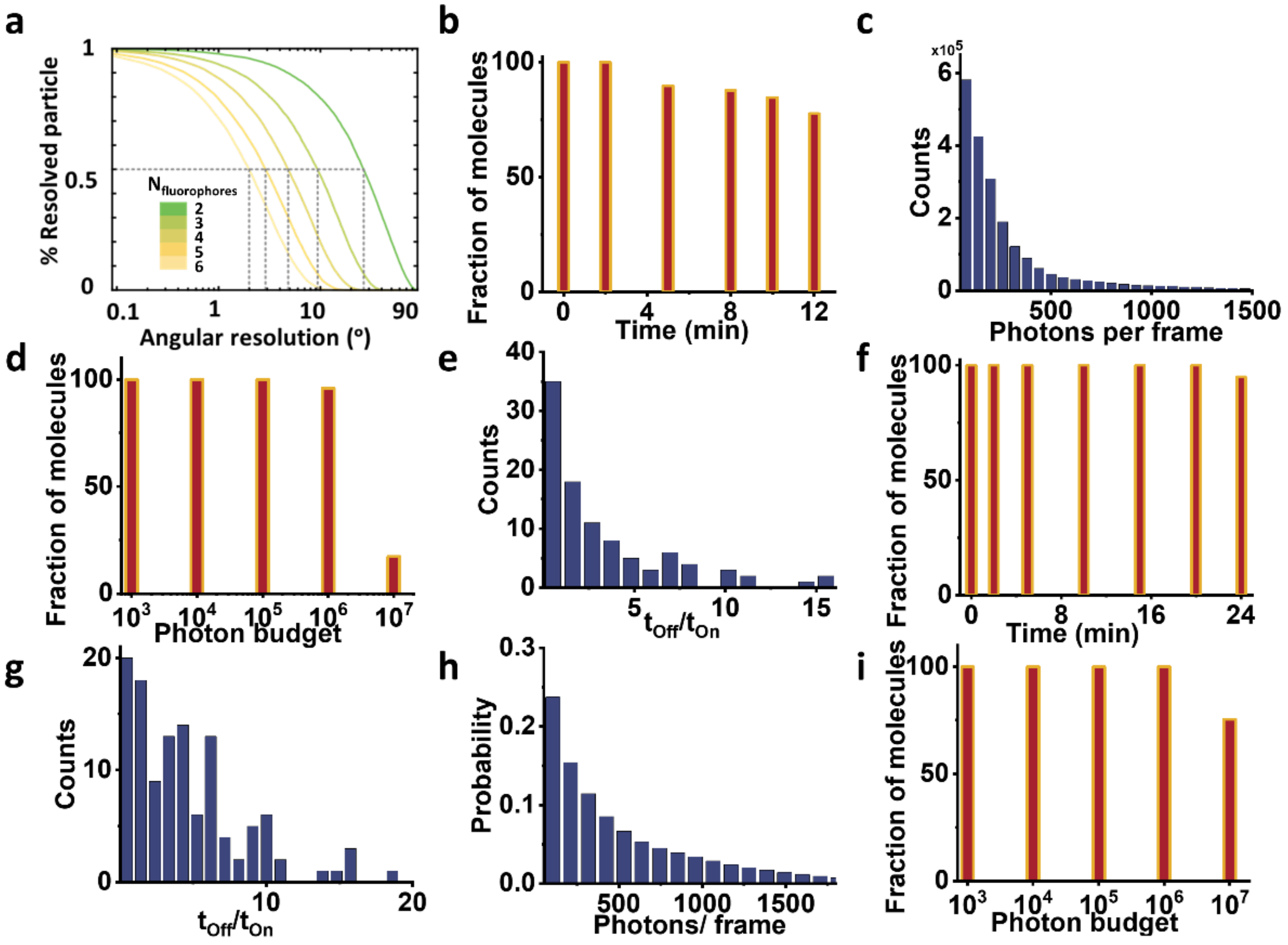


**Figure S2.**

(a) The fraction of particles for which all fluorophores can be identified by the analysis of their polarization time-trace is shown as a function of the angular resolution of the fit. In order to resolve 50% of a hexamer sample with 6 fluorophores, an angular resolution of 2° is required. However, to resolve a partially labeled hexamer with 3 fluorophores only requires a resolution of 10°. For this simulation we assumed random dipole angles and accounted for the mapping into the [0°, 90°] interval.

(b-e) Photo-physics characterization of NTA-ATTO647N attached to human PCNA protein imaged at 4 K.

(b) Fluorophore stability histogram shows the fraction of molecules which survived until a given time of data acquisition. Most molecules survived until the end of the measurement, allowing improved photon collection and higher localization precision.

(c) Histogram of the mean number of photons per frame. On average we obtain ~ 260 photons per frame.

(d) Photon budget histogram plotted as a fraction of molecules versus total number of photons collected from 50,000 frames. Around 17 % of the molecules emitted at least $10^7$ photons. However, this number is mostly limited by the shorter recording time in these measurements and can be increased further if necessary.

(e) The photo-switching behavior of the fluorophores is characterized by the dwell times in the off- and on-state. The ratio of their average duration indicates long-lived off-state facilitate localizing single emitters via polarization-trace fitting.

(f-i) Photo-physics characterization of ATTO647N-maleimide attached ClpB protein imaged at 4 K.

(f) The fluorophore stability histogram demonstrates a high survival fraction of molecules.
(g) Histogram of the off-on ratio.
(h) The mean number of photons per frame shows a two-fold increase compared to the PCNA trimer with an average of ~510 photons.
(i) Photon budget histogram plotted as a fraction of molecules versus total number of photons collected from 100,000 frames. Around 75 % of the molecules emitted at least $10^7$ photons.

### S3 - Example of complete intensity and polarization trace processing for fluorophore localization

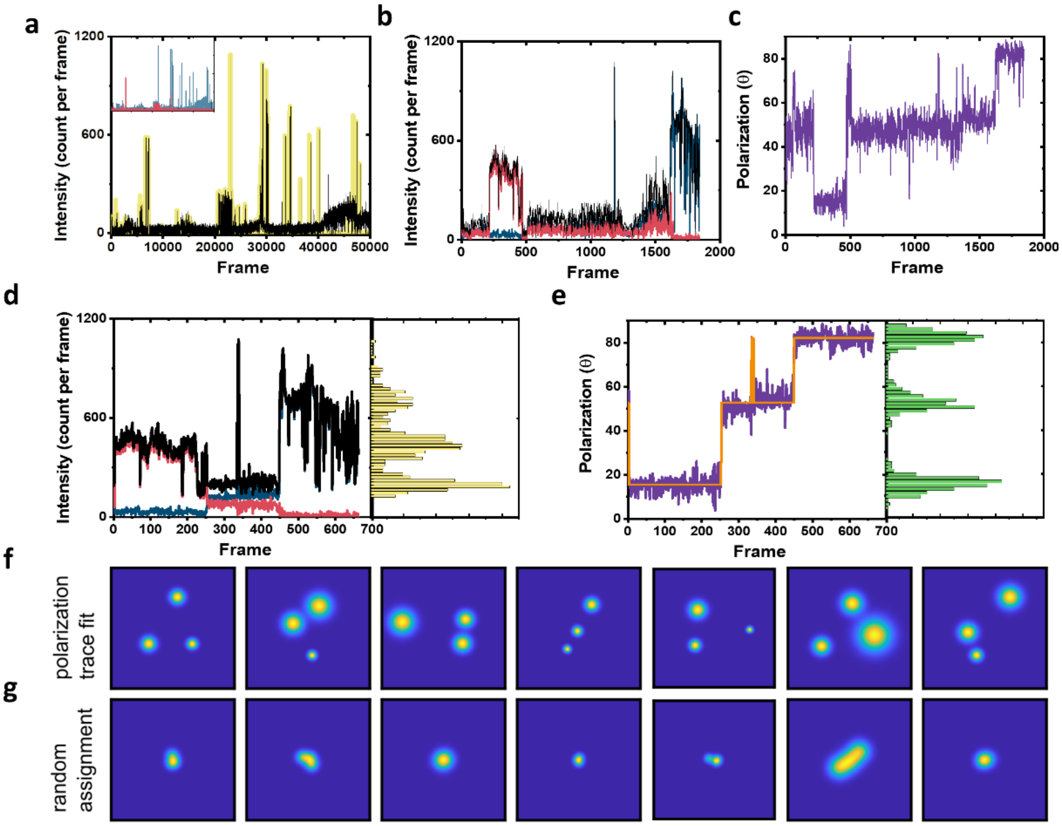


**Figure S3.**

(a-e) Example of complete signal processing calculated from the same single molecule.

(a) Raw total photon counts registered from the two channels (black line). Yellow highlights show the regions which were determined as on events and taken for further analysis. The inset shows the photon signal in each channel separately shown in blue and red.

(b) Intensity time trace of the on events highlighted in (a). The blue and red lines represent the intensity of the two polarization channels, and the black line is the total signal.

(c) Polarization trace (purple color) of the signal calculated from (b).

(d) The signal trace after filtering based on photon counts per frame (frames below 100 photons were excluded) with the yellow histogram on the side showing the photon distribution over the whole measurement. The color code is the same as in (b).

(e) The polarization trace (purple color) as calculated from intensity signal shown in (d) and used for fluorophore assignment and further analysis. The green histogram on the side shows the polarization distribution. The Orange solid line is the fit of the polarization trace as calculated using DISC algorithm, which combines unsupervised statistical learning tools with change-point detection in a model-independent manner (1). The fit clearly show 3 separate polarization states, which indicate the fluorophores attached to this molecule. Next, we take the assignment of the polarization trace to assign and localize the coordinate of each fluorophore separately.

(f) Example of 2D maps of the fluorophore localization as obtained for each fluorophore based on fitting the polarization trace in (e).

(g) In order to validate the assignment of the detected photons originating from a diffraction-limited spot to a specific fluorophore, we performed random assignment of frames. After averaging these randomly assigned localizations, the super-resolved image shows single spots in the center. In comparison, the polarization time trace fit accurately resolves the 3 positions in (f).

### S4 - Complete data set of the PCNA trimer used for 3D reconstruction

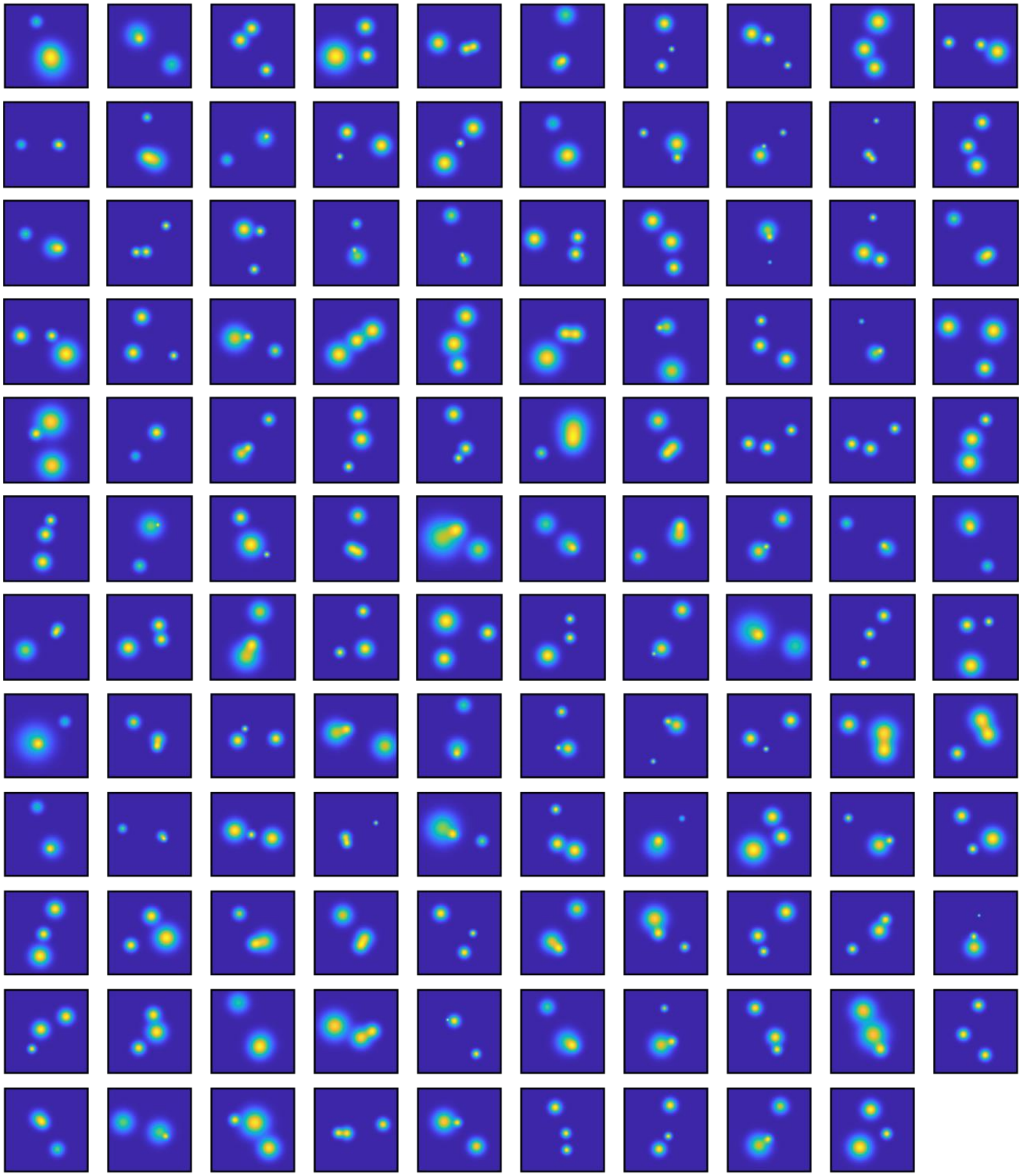

**Figure S4.** All 2D maps as obtained from polarization trace fitting with a three-state model. Particles were filtered based on localization precision below 3 nm. The image size is 120 x 120 pixel at 0.15 nm/pixel.

### S5 - ClpB labeling and 3D reconstruction of classified 2D images

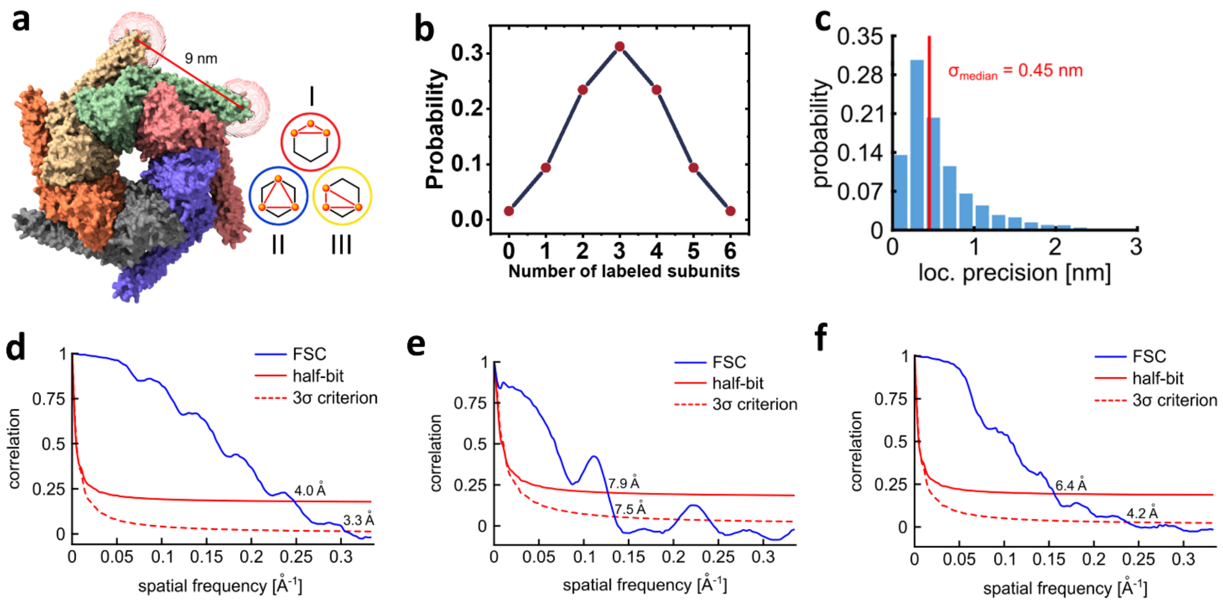


**Figure S5.** (a) We modeled the dye positions using the method described previously in Fig. 2b. In particular, we calculated the accessible volume of the dye-linker attached to the sulfur group of a cysteine amino acid at position 428. We used dye parameters for ATTO647N maleimide as provided in Ref. (2). We used the center of mass of the obtained cloud coordinates to calculate the expected pair-wise distances between fluorophores and obtained a distance of 9 nm between two adjacent subunits. Given the planar hexameric geometry, the other possible pair-wise distances are 15.5 nm and 18 nm. Inset shows the three possible configurations, classes I, II and III obtained from particles labeled with three fluorophores.

(b) The fraction of labeled protomers in ClpB molecules was calculated based on the binomial distribution. Here, we assume 50% labeling efficiency and therefore expect that 1/3 of proteins will carry 3 fluorophores.

(c) Histogram of localization precisions with a median of 0.45 nm obtained from the hexamer data.

(d-f) FSC of the 3D volumes obtained for each class separately. For each class, 2D images were divided equally into two halves randomly. Then 3D reconstitution was calculated for each half data set separately, using a low-pass filtered model as an initial guess. The correlation between the two 3D reconstitution is calculated over shells of the Fourier transform and plotted as a function of spatial frequency (blue curves). The spatial resolution of the 3D reconstitution is then evaluated using half-bit criteria (red solid line) and 3σ criterion (red dashed line) as described in details in (3).

(d) The FSC of class I yields a resolution of 4.0 Å.

(e) The FSC of class II yields a resolution of 7.9 Å.

(f) The FSC of class III yields a resolution of 6.4 Å.

### S6 - Data set of the ClpB hexamer protein used for classification

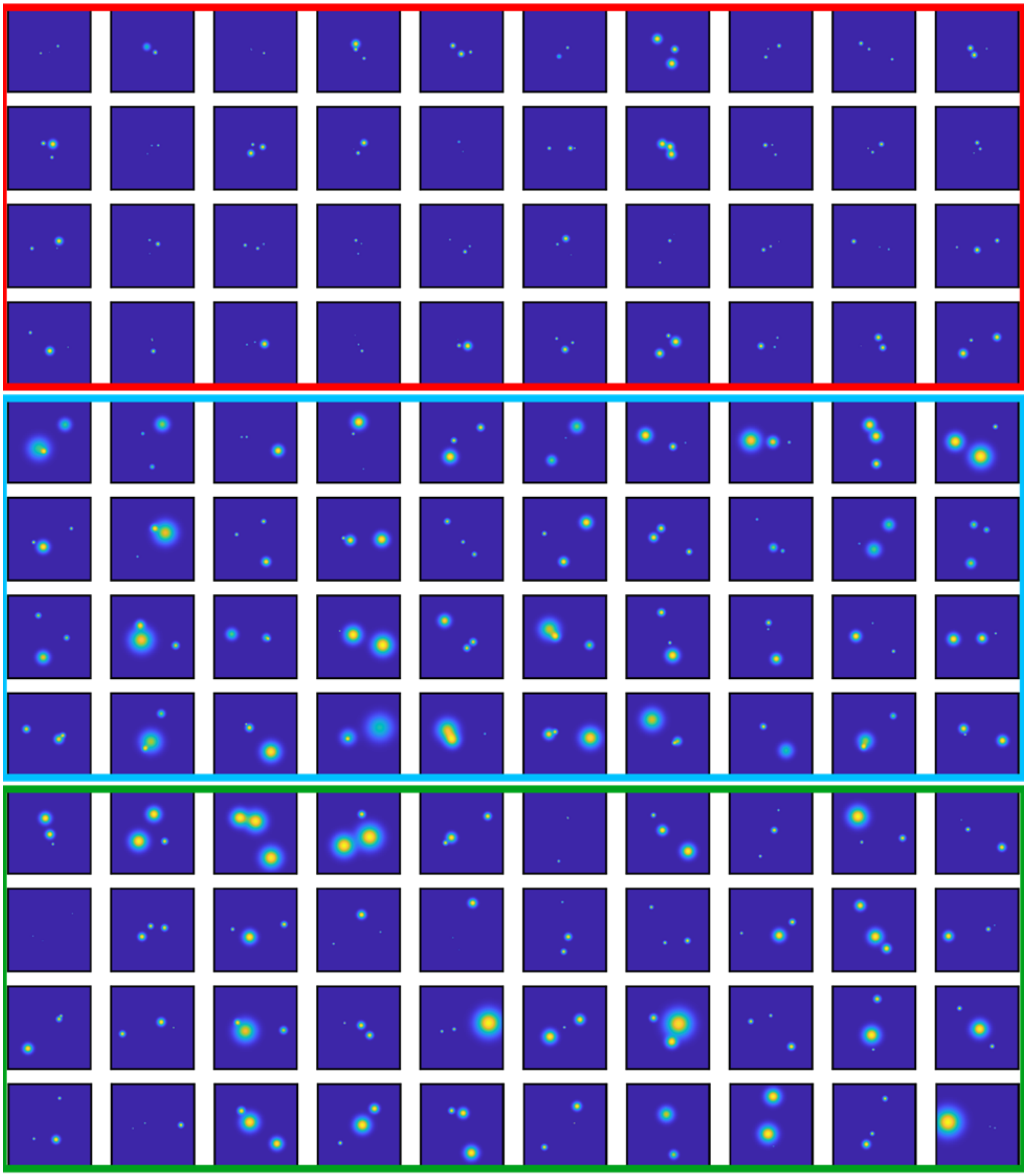

**Figure S6.** Some examples of the resolved 2D maps as obtained from polarization trace fitting with a three-state model. Particles were filtered based on a localization precision below 3 nm and classified (red = class I, blue= class II, green = class III). The image size is 200 x 200 pixel at 0.15 nm/pixel.

### S7 - 3D reconstitution validation of ClpB classification

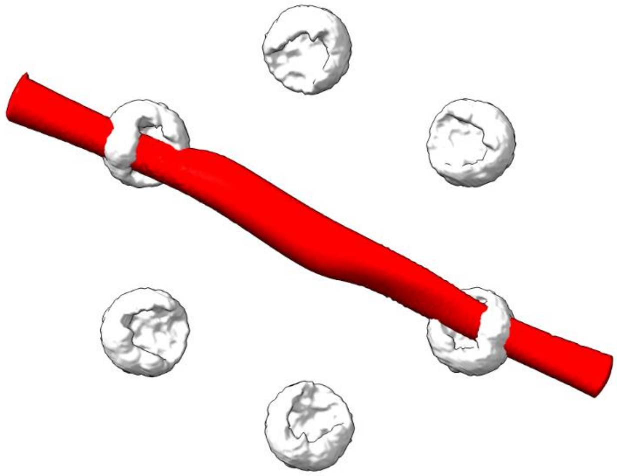

**Figure S7.** Validation of the particle classification of the hexamer protein ClpB. Here, we took all the unclassified 2D images with correlation scores below 0.8, and calculated the 3D reconstitution as described previously in method section of main text. The red color shape in the image shows no valid structure and could not be fitted the white clouds which represent the accessible volumes of the fluorophores as calculated for each protomer of the hexamer protein.

### S8 - Correlative polarCOLD and cryoEM on the single particle level

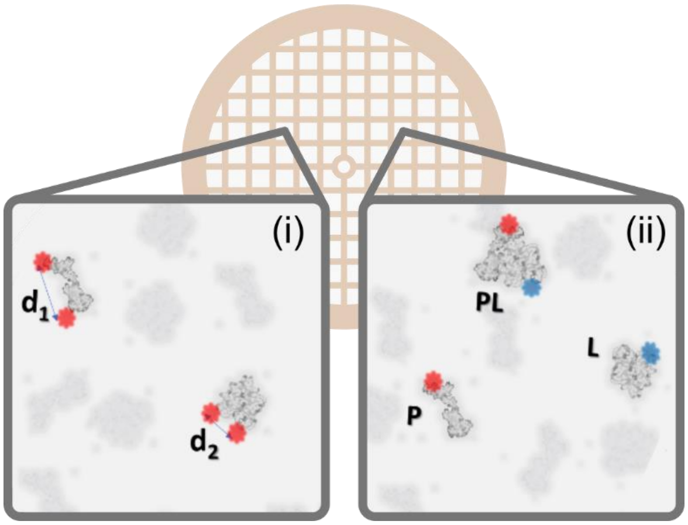


**Figure S8.** By measuring the polarization signal of multiple fluorophores one can classify the particles in a low-contrast EM sample based on distances and orientation. In the exemplary case on the left (i) molecules are identified based on intermolecular or intramolecular distances, indicated by $d_1$ and $d_2$. The case on the right (ii) illustrates the sorting and sizing of interacting molecules using nanometer distance measurement with multi-color polarCOLD, where Apo proteins (P) and ligand (L) are labeled with different fluorophores, such that the resulting complex (PL) exhibits fluorescence at both wavelengths.

**Table S1** | Overview of the recorded data and the experimental yield

| Protein | Human PCNA | ClpB | | |
|---|---|---|---|---|
| Total number of particles | 7593 | 23765 | | |
| Particles identified with three polarization states | 2543 | 6259 | | |
| Particles per class (Classification) | - | C1 | C2 | C3 |
| | | 3806 | 407 | 2046 |
| Particles used for 3D reconstitution | 119 | 232 | 100 | 135 |
| Yield | 4,67 % | 7,46 % | | |

### References

1. D. S. White, M. P. Goldschen-Ohm, R. H. Goldsmith, B. Chanda, Top-down machine learning approach for high-throughput single-molecule analysis. *eLife* **9** (2020).
2. S. Kalinin *et al.*, A toolkit and benchmark study for FRET-restrained high-precision structural modeling. *Nat Methods* **9**, 1218-1225 (2012).
3. M. Van Heel, M. Schatz, Fourier shell correlation threshold criteria. *Journal of Structural Biology* **151**, 250-262 (2005).
